# Infection increases activity via *Toll* dependent and independent mechanisms in *Drosophila melanogaster*

**DOI:** 10.1101/2021.08.24.457493

**Authors:** Crystal M. Vincent, Esteban J. Beckwith, William H. Pearson, Katrin Kierdorf, Giorgio Gilestro, Marc S. Dionne

## Abstract

Host behavioural changes are among the most apparent effects of infection. ‘Sickness behaviour’ can involve a variety of symptoms, including anorexia, depression, and changed activity levels. Here we use a real-time tracking and behavioural profiling platform to show that, in *Drosophila melanogaster*, many systemic bacterial infections cause significant increases in physical activity, and that the extent of this activity increase is a predictor of survival time in several lethal infections. Using various bacteria and *D. melanogaster* immune and activity mutants, we show that increased activity is driven by at least two different mechanisms. Increased activity after infection with *Micrococcus luteus*, a Gram-positive bacterium rapidly cleared by the immune response, strictly requires the *Toll* ligand *spätzle* and Toll-pathway activity in the fat body and the brain. In contrast, increased activity after infection with *Francisella novicida*, a Gram-negative bacterium that cannot be cleared by the immune response, is entirely independent of either *spätzle* or the parallel IMD pathway. The existence of multiple signalling mechanisms by which bacterial infections drive increases in physical activity implies that this effect may be an important aspect of the host response.

## Introduction

Some of the most apparent effects of infection are the sickness behaviours of the host. A variety of infection-induced behavioural changes have been documented; in vertebrates, these commonly include anorexia, lethargy, and social withdrawal (Dantzer, 2001; Shattuck and Muehlenbein, 2015; Stockmaier et al., 2021). In insects, a partially-overlapping set of changes have been described, including anorexia and foraging changes, behavioural fevers, and changes in oviposition (Anderson et al., 2013; Masuzzo et al., 2019; Stahlschmidt and Adamo, 2013; Surendran et al., 2017; Vale and Jardine, 2017). These changes in behaviour can in some cases facilitate immune function, either in terms of pathogen clearance or host survival; in other cases, the pathogen appears to benefit; and in some cases, there is no obvious beneficiary, and the observed behavioural change may be a non-selected consequence of the interaction of two or more complex physiological systems. In all of these cases, however, infection behaviours have a strong effect on the well-being of the host, whether or not this effect is ultimately manifested as a difference in infection outcome.

Whilst sickness behaviours are often described as being part of the host response to infection, several studies have shown that behavioural changes during infection can also be the result of pathogen manipulation of host biology, rather than the host responding to a pathogen threat, *per se* (Adamo and Webster, 2013; Berdoy et al., 2000; Heil, 2016; Klein, 2003; Lafferty and Shaw, 2013). The difference between a host response and parasite manipulation is not simply a matter of semantics as the two can predict opposing evolutionary trajectories and infection outcomes (Hart, 1988; Johnson, 2002; Klein, 2003). If hosts change their behaviour in response to the physiological stresses associated with infection, we assume that said behaviour will be of benefit to the host, usually by reducing pathology (Ayres and Schneider, 2009; Kuo and Williams, 2014; Vincent and Bertram, 2010; Wang et al., 2016). In contrast, when pathogens manipulate host behaviour, we assume it serves the function of increasing pathogen fitness, often via enhanced transmission (Andersen et al., 2009; Berdoy et al., 2000; Biron et al., 2006; Shaw et al., 2009; Thomas et al., 2005, 2002; Webster et al., 1994).

Changes in sleep and activity are some of the most common behavioural manifestations of infection, seen in vertebrates and invertebrates (Besedovsky et al., 2019; Kuo et al., 2010; Shirasu-Hiza et al., 2007). The extensive crosstalk between sleep and immunity has led to many suppositions regarding the value of sleep in maintaining a robust immune response and health in the face of infection (Besedovsky et al., 2019; Imeri and Opp, 2009; Kuo et al., 2010; Majde and Krueger, 2005; Opp, 2009). However, despite investigations of the interplay between sleep and infection in insects, there remain inconsistencies in whether sleep (or activity) is induced or inhibited during infection, and what effect these changes have on infection pathology (Arnold et al., 2013; Kuo et al., 2010; Kuo and Williams, 2014; Mallon et al., 2014; Shirasu-Hiza et al., 2007; Siva-Jothy and Vale, 2019). Whilst some of this incongruity may result from the fact that in these studies flies were injected at different times of the day (Lee and Edery, 2008), they could also be caused by differences between pathogens used and therefore disparate activation of immune factors (Hoffmann, 2003; Lemaitre et al., 1997, 1995; Tanji and Ip, 2005; Wang and Ligoxygakis, 2006). The consequences of infection-induced changes in sleep and activity are thus multifaceted and the effects of infection will depend on interaction between host immune and nervous systems and the pathogen itself on multiple levels.

Using the real-time tracking and behavioural profiling platform, the ethoscope (Geissmann et al., 2017), we test an array of various bacteria and *D. melanogaster* immune and activity mutants, to determine whether pathogen recognition and immune pathway activation contribute to the increase in activity observed during infection.

## Results

### Bacterial infection leads to a marked increase in locomotor activity

We began by exploring the effects of *Francisella novicida* on physical activity in *Drosophila melanogaster*; a Gram-negative bacterium that propagates both intra- and extra-cellularly in *D. melanogaster*, ultimately resulting in host death after four days (Moule et al., 2010; Vincent et al., 2020; Vonkavaara et al., 2008). This infection is particularly tractable for behavioural studies because it presents an infection course in excess of three days (two days in some immune mutants), allowing ample time for activity monitoring; near-synchronous mortality; and strong immune activation, allowing identification of effects of immune activation on activity (Moule et al., 2010; Vincent et al., 2020). We found that flies infected with *F. novicida* spent 10% and 9% more time moving than mock injected and uninfected controls, respectively (Figure 1A). This increase in activity intensified over the course of infection. Further partitioning of activity data found that while there were subtle increases in micro-movements such as grooming and feeding (Geissmann et al., 2019, 2017) (Figure 1B), the observed increase in movement was primarily the result of increased time spent walking; these flies also spent 12% and 11% less time sleeping (Figure 1C, D; Figure 1 – figure supplement 1A-C). Importantly, despite spending proportionately more time active, infected flies did not cover a greater total distance, indicating that the intensity of their activity was unaltered by infection (Figure 1 – figure supplement 1D, E). Activity on the first day of infection was predictive of lifespan, with more active flies exhibiting increased lifespan (Figure 1E). In addition, there was a positive correlation between total activity and survival (Figure 1 – figure supplement 1F); we used day one activity as a predictor because previous work has found immune activity to be strongest at this time (De Gregorio et al., 2002a; Lemaitre et al., 1997; Schlamp et al., 2021) and thus would be an appropriate metric in looking at infections of shorter duration. Furthermore, day 1 activity levels were positively correlated with total activity levels (Figure 1 – figure supplement 1G), giving us confidence that activity on day 1 is representative of total activity levels in assessing infections of longer duration.

**Figure 1.**
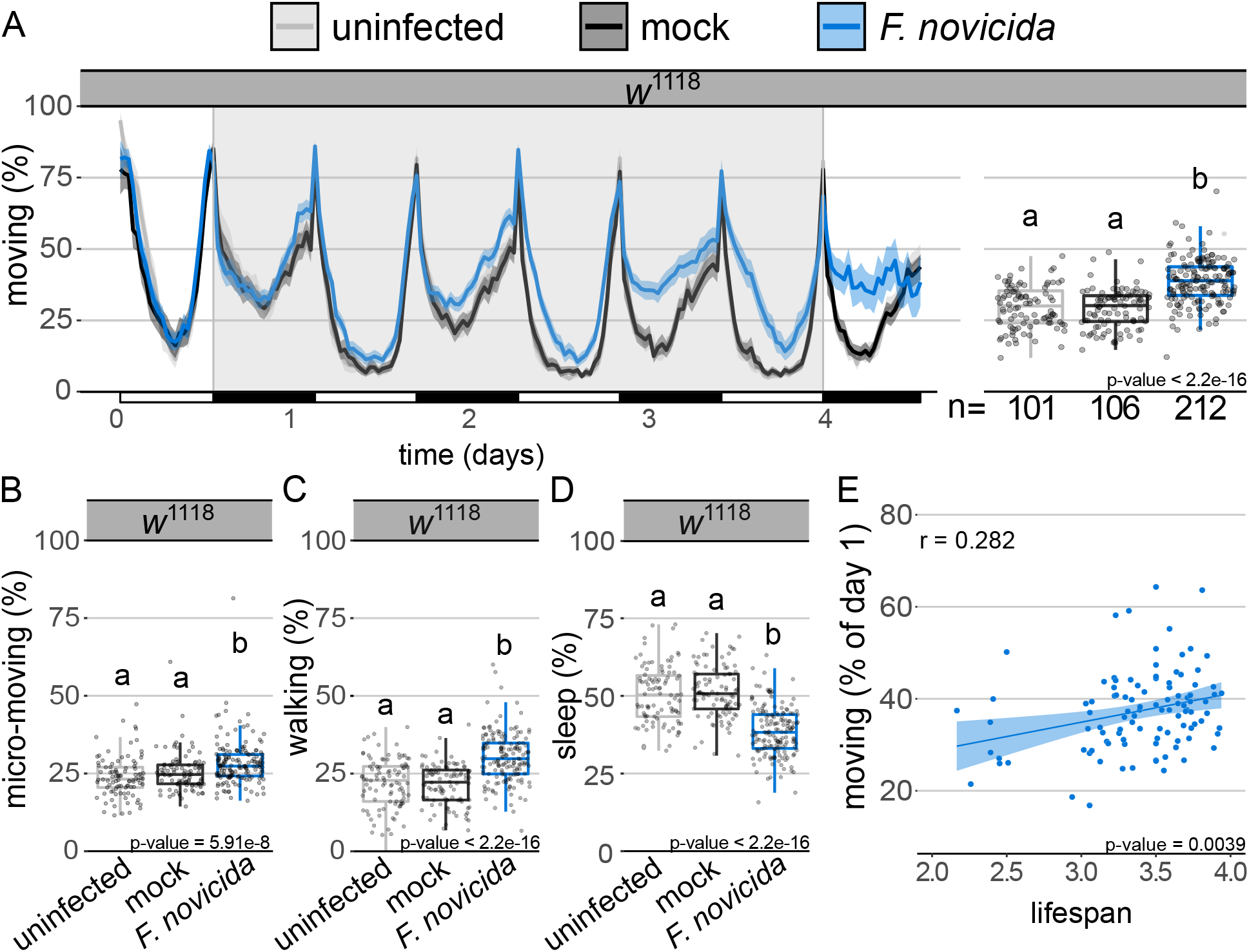
Infection with *Francisella novicida* leads to increased locomotor activity. **(A)** Ethogram showing percentage of wild-type males moving over time. Alternating white and black horizontal bar along the x-axis indicates day (12h light) and night (12h dark) cycles, respectively. Uninfected and mock controls are represented by grey and black tracings, respectively. Infected flies are in blue. Shaded areas surrounding solid lines represent the 95% confidence intervals. Flies were injected within two hours of the beginning of their light cycle (t = 0). Background area highlighted in grey indicates the time for which data were analysed as represented in adjoining boxplot. Boxplots showing percentage of infected wild-type males **(B)** engaging in micro-movements (e.g. feeding and grooming), **(C)** walking and **(D)** sleeping. Markers indicate individual data points. Horizontal bar within each box represents the median. The bottom and top lines of the box represent the 1^st^ and 3^rd^ quartiles, respectively. Whiskers represent the smallest value between: highest and lowest values or 1.5x the interquartile range. Boxes without common letters are significantly different. Sample sizes (n) are indicated under the boxplots. Plots throughout have identical formatting, therefore a full description of ethogram and boxplot features is omitted in subsequent legends. *Francisella novicida* infected animals moved significantly more than both the uninfected and mock controls (Kruskal-Wallis chi-square = 99.206, df = 2, n = 419, p = 2.2e-16; Dunn’s *post hoc*: mock|*F. novicida* = 1.2e-17, mock|uninfected = 0.41, uninfected|*F. novicida* = 1.4e-14). Infected flies **engaged in more micro-movements** (Kruskal-Wallis chi-square = 33.287, df = 2, n = 419, p = 5.9e-08; Dunn’s *post hoc*: mock|*F. novicida* = 9.5e-05, **walked more** (Kruskal-Wallis chi-square = 88.383, df = 2, n = 419, p = 2.2e-16; Dunn’s *post hoc*: mock|*F. novicida* = 1.02e-15, mock |uninfected = 0.48, uninfected|*F*. *novicida* = 2.7e-13), mock |uninfected = 0.23, uninfected|*F*. *novicida* = 2.7e-07), and **spent less time sleeping** (Kruskal-Wallis chi-square = 99.206, df = 2, n = 419, p = 2.2e-16; Dunn’s *post hoc*: mock|*F. novicida* = 1.2e-17, mock|uninfected = 0.41, uninfected|*F*. *novicida* = 1.4e-14) than both the mock and uninfected controls. **(E)** Activity level within the first day of *F. novicida* infection was positively correlated with survival (Pearson’s correlation, *r* = 0.282; t = 2.96, df = 101, p = 3.9e-03). Data from multiple replicates are shown.

To test whether greater activity following infection was specific to this infection or a general consequence of immune activation, we infected wild-type flies with a phylogenetically and pathogenically diverse panel of bacteria. We found that three of the five bacteria examined, *Micrococcus luteus, Listeria monocytogenes*, and *Staphylococcus aureus*, induced increased activity (Figure 2A-C; Figure 2 – figure supplement 1). Thus, including *F. novicida*, the four bacteria able to drive hyperactivity include Gram-positives and Gram-negatives, as well as microbes killed efficiently by the immune response and those able to survive within and outside of host cells (Hanson et al., 2019; Moule et al., 2010; Nehme et al., 2011; Vincent et al., 2020). As we found in *F. novicida*, activity on the first day of infection was predictive of lifespan, with more active flies exhibiting increased lifespan during *L. monocytogenes* infection, but decreased lifespan during infection with *S. aureus* (Figure 2D, E). Next, we screened a selection of immune, locomotor and circadian mutants of *D. melanogaster* for activity levels during *F. novicida* infection and observed increased locomotor activity in all mutants tested (Figure 2 – figure supplement 2; Figure 2 – figure supplement table 1). Thus, we concluded that the effect of infection on locomotor behaviour is a widespread phenomenon and may represent a complex trait emerging as the result of the induction of multiple molecular pathways.

**Figure 2.**
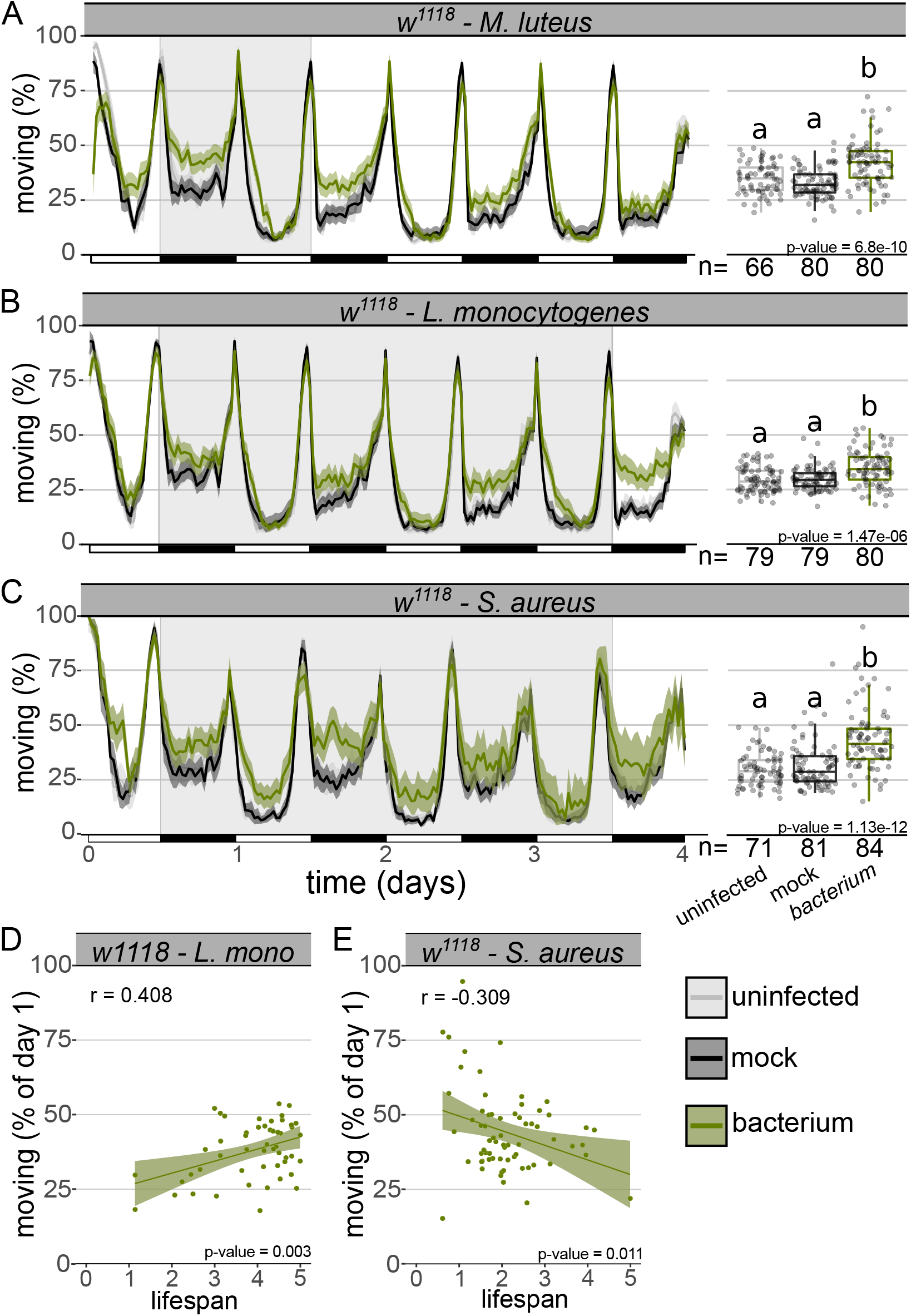
Infection with multiple bacteria leads to increased activity in wild-type flies. Ethogram showing percentage of wild-type flies moving over time during infection with **(A)** *Micrococcus luteus* **(B)** *Listeria monocytogenes* and **(C)** *Staphylococcus aureus*. Infected flies moved significantly more than both the uninfected and mock controls (***M. luteus***: Kruskal-Wallis chi-square = 42.22, df = 2, n = 226, p = 6.8e-10; Dunn’s *post hoc*: mock|*M. luteus* = 7.02e-10, mock|uninfected = 0.07, uninfected|*M. luteus* = 3.0e-05; ***L. monocytogenes***: Kruskal-Wallis chi-square = 26.859, df = 2, n = 238, p = 1.5e-06; Dunn’s *post hoc*: mock|*L.monocytogenes* = 3.05e-05, mock|uninfected = 0.68, uninfected|*L.monocytogenes* = 8.7e-06; ***S. aureus***: Kruskal-Wallis chi-square = 55.016, df = 2, n = 236, p = 1.1e-12; Dunn’s *post hoc*: mock|*S. aureus* = 1.2e-09, mock|uninfected = 0.49, uninfected|*S. aureus* = 1.1e-10). Activity level within the first day of **(D)** *L. monocytogenes* and **(E)** *S. aureus* infection was positively correlated with survival (*L. monocytogenes*: Pearson’s correlation, *r* = 0.408; t = 3.13, df = 49, p = 2.9e-03; *S. aureus*: Pearson’s correlation, *r* = − 0.309; t = − 2.61, df = 64, p = 0.0114). Data from multiple replicates are shown. Behavioural assays were performed at least twice, data from all replicates are shown.

### Increased activity is not a moribund behaviour and is affected by immune activation

We became particularly interested in the change in locomotor activity observed during infection with *M. luteus* because, unlike the other bacteria examined, increased activity following injection with *M. luteus* was transient (Figure 2A). That flies spend more time active during *M. luteus* infection is particularly important because it demonstrates that infection-induced activity is not a moribund behaviour: *M. luteus* infection is cleared by the immune response and flies are not killed by this infection over the following four days (Figure 2 – figure supplement 3); this contrasts with *F. novicida*, *L. monocytogenes* and *S. aureus*, which kill more than half of all infected flies within four days (Figure 2 – figure supplement 3).

In *M. luteus* and *F. novicida*-infected flies, infection-dependent increases in activity were roughly correlated with bacterial load. In flies infected with *M. luteus*, activity increased during the early stages of infection when bacterial numbers were high and declined once bacteria had been cleared. In *F. novicida*-infected flies, activity increased in parallel with the bacterial load (Figure 2 – figure supplement 3). This parallel between these infections prompted us to test whether bacterial detection by immune pathways and the subsequent signalling drove increased activity. Previous work found that the NFκB transcription factor RELISH which plays a vital role in *D. melanogaster’s* immune response, is required for infection-induced sleep (Kuo et al., 2010). We infected flies lacking the Toll and immune deficiency (IMD) pathways, the primary microbe-detection pathways in *D. melanogaster* (De Gregorio et al., 2002b; Lau et al., 2003; Lemaitre et al., 1995; Tanji and Ip, 2005). We found that ablation of IMD (*imd*^10191^) and Toll signalling (*spz*^eGFP^) had disparate effects during infection. Activity during *M. luteus* infection was unaffected in *imd* mutants, but no increase in activity was observed following *M. luteus* infection in *spz* mutants (Figure 3A, B), in keeping with the fact that *M. luteus* is primarily an agonist of the Toll pathway (Irving et al., 2001; Lemaitre et al., 1997; Rutschmann et al., 2002). To confirm this finding, we repeated this experiment using flies carrying a different *spz* allele (Kenmoku et al., 2017) and found the same result (Figure 3C).

**Figure 3.**
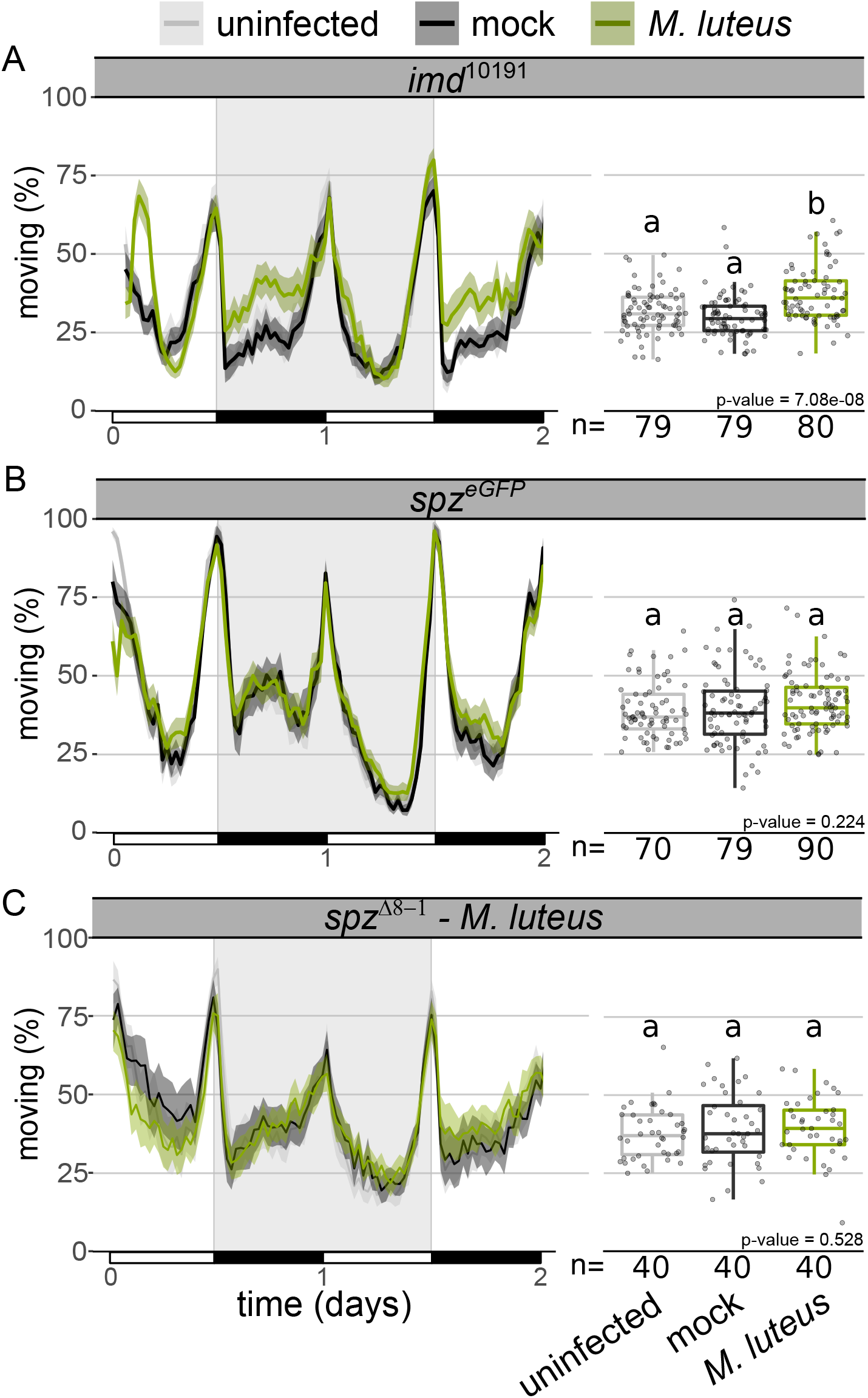
Toll signalling mutants do not increase activity during *M. luteus* infection. Ethogram showing percentage of **(A)** *imd*^10191^, **(B)** *spz*^eGFP^ and **(C)** *spz*^Δ8-1^ flies moving over time. *Micrococcus luteus* infected *imd*^10191^ flies –but not *spz* ^eGFP^ –moved significantly more than both the uninfected and mock controls (*imd*^10191^: Kruskal-Wallis chi-square = 32.93, df = 2, n = 238, p = 7.1e-08; Dunn’s *post hoc*: mock|*M. luteus* = 6.9e-08, mock|uninfected = 0.1, uninfected| *M. luteus* = 1.07e-04; *spz* ^eGFP^: Kruskal-Wallis chi-square = 2.99, df = 2, n = 239, p = 0.22; ***spz*** ^Δ8-1^: Kruskal-Wallis chi-square = 1.28, df = 2, n = 120, p = 0.528). Data from multiple replicates are shown.

The absence of increased activity in *M. luteus*-infected Toll mutants indicates that Toll signalling is required for hyperactivity during *M. luteus* infection. However, mutation of either *imd* or *spz*, as well as the combination of the two (*imd*^10191^; *spz*^eGFP^), did not affect the increase in activity caused by *F. novicida* (Figure 3 – figure supplement 1), supporting our previous results with *Tak1* mutants (Figure 2 – figure supplement 2; Figure 2 – figure supplement table 1). These findings demonstrate that in *F. novicida* infections, the activity phenotype is independent of *Toll* and *imd* pathways. The dependence on Toll for increased activity in *M. luteus* but not *F. novicida* infections indicates that infection induces activity via different signalling pathways during these infections.

### Infection causes temporally-specific metabolic dysregulation, but infection-induced activity is not a response to starvation

Infection with *M. luteus* inhibits insulin signalling, as evidenced through a reduction in phosphorylated AKT, and this metabolic shift is concomitant to a reduction in triglyceride levels (DiAngelo et al., 2009). Similarly, infection with *F. novicida* leads to triglyceride loss as well as hyperglycaemia and reduced levels of glycogen (Vincent et al., 2020). The interplay between immune and metabolic signalling pathways is thought to be indicative of the metabolic burden associated with infection and the need to redistribute available resources (Clark et al., 2013; DiAngelo et al., 2009; Dionne et al., 2006). We therefore surmised that the metabolic shifts observed during *M. luteus* and *F. novicida* infection could play a role in infection-induced activity, despite the difference in the requirement of Toll signalling, and sought to determine whether *F. novicida* and *M. luteus* infections led to similar metabolic phenotypes in flies.

We tested whether infection with *F. novicida* inhibits insulin signalling as has been previously reported for *M. luteus* infection (DiAngelo et al., 2009). We found that infection with *F. novicida* inhibits insulin signalling as determined through the observance of lower levels of phosphorylated-AKT during late infection (72-80h post-injection; Figure 4A); these flies were also hyperglycaemic and exhibited depleted triglyceride and glycogen stores (Figure 4B). Similarly, late in infection with *M. luteus*, flies had lower triglyceride and glycogen levels but no change in circulating sugars. During early infection (24-30h post-injection), we observed hypoglycaemia with *F. novicida*, but not *M. luteus*, decreased triglycerides with *M. luteus*, but not *F. novicida*, and low levels of glycogen in both infections (Figure 4B). These results confirm previous work showing that bacterial infection can lead to metabolic pathology including hyperglycaemia and loss of triglyceride and glycogen stores and that these effects are often limited to specific times over the course of an infection (Chambers et al., 2012; Dionne et al., 2006; Vincent et al., 2020).

**Figure 4.**
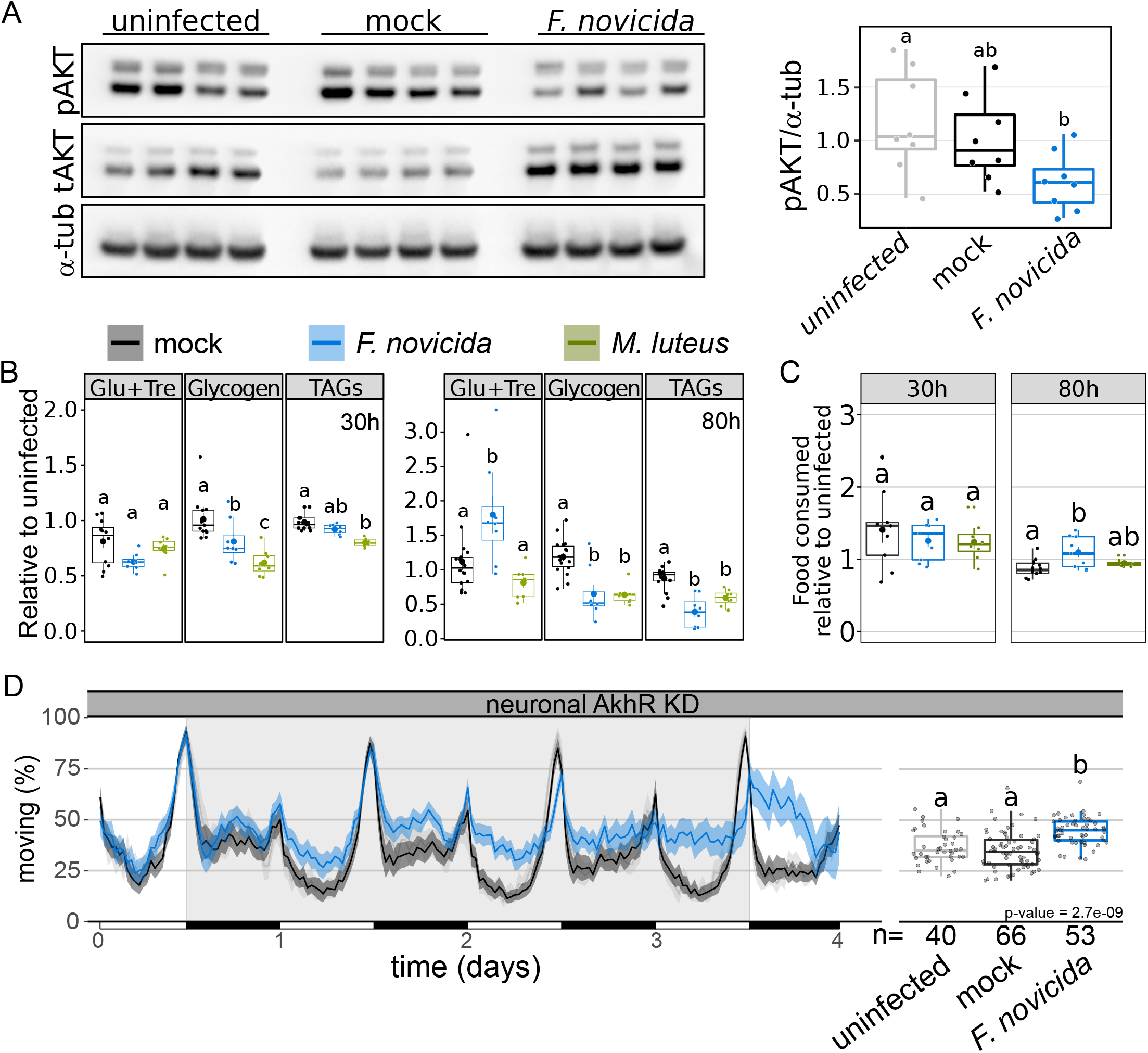
*Micrococcus luteus* and *Francisella novicida* infection similarly disrupt host metabolism. **(A)** Western blot of phosphorylated AKT (Ser505) during *F. novicida* infection in wild-type flies. Total AKT and tubulin levels for each sample also shown. Boxplot shows quantification of pAKT relative to α-tubulin using data from both repeats of the experiment. **(B)** Levels of circulating and stored glycogen and triglyceride (TAG) and feeding activity **(C)** during early (30h) and late (80h) infection. Mock controls are indicated in grey, whilst *F. novicida* and *M. luteus* injections are indicated in blue and green, respectively. Data are plotted relative to the mean of uninfected controls. **Metabolism 30h post infection**: There was a significant effect of infection, such that glucose levels were significantly lower in *F. novicida*-infected flies (AOV: df = 2, n = 28, F = 3.449, p = 0.048; Tukey’s HSD: mock|*F. novicida* = 0.038, mock |*M. luteus* = 0.62, *M. luteus*|*F. novicida* = 0.29). Infection had a significant effect on glycogen stores which were reduced in both infections (AOV: df = 2, n = 29, F = 12.112, p = 2.1e-04; Tukey’s HSD: mock|*F. novicida* = 0.049, mock |*M. luteus* = 1.4e-04, *M. luteus*|*F. novicida* = 0.102), but only *M. luteus-*infected flies had a significant reduction in triglycerides (AOV: df = 2, n = 27, F = 18.763, p = 1.5e-05; Tukey’s HSD: mock|*F. novicida* = 0.073, mock|*M. luteus* = 8.8e-06, *M. luteus*|*F. novicida* = 3.5e-03). **Metabolism 80h post infection**: There was a significant effect of infection, such that glucose levels were significantly higher in *F. novicida*-infected flies (AOV: df = 2, n = 31, F = 6.883, p = 3.5e-03; Tukey’s HSD: mock|*F. novicida* = 0.018, mock|*M. luteus* = 0.41, *M. luteus*|*F. novicida* = 3.4e-03). Infection led to a significant reduction of both glycogen (AOV: df = 2, n = 31, F = 17.315, p = 1e-05; Tukey’s HSD: mock|*F. novicida* = 1.6e-04, mock |*M. luteus* = 1.1e-04, *M. luteus*|*F. novicida* = 0.99), and triglycerides (AOV: df =2, n = 28, F = 21.622, p = 2.5e-06; Tukey’s HSD: mock|*F. novicida* = 2.3e-06, mock|*M. luteus* = 2.7e-07, *M. luteus*|*F. novicida* = 0.066). **Feeding**: Neither infection affected feeding within **30h** of injection (AOV: df =2, n = 26, F = 1.117, p = 0.35), but **80h** post-injection, *F. novicida-*infected flies fed significantly more than mock controls but not more than *M. luteus*-infected flies (AOV: df =2, n = 29, F = 7.289, p = 4.2e-03; Tukey’s HSD: mock|*F. novicida* = 9.2e-03, mock|*M. luteus* = 0.58, *M. luteus*|*F. novicida* = 0.082). **(D)** Ethogram showing percentage of flies moving over time. Neuronal KD of adipokinetic hormone did not eliminate *F. novicida* infection-induced activity, suggesting that this phenotype is not resulting from starvation (Kruskal-Wallis chi-square = 39.461, df =2, n = 159, p = 2.7e-09; Dunn’s *post hoc*: mock|*F. novicida* = 4.3e-09, mock|uninfected = 0.35, uninfected|*F. novicida* = 1.4e-05). Data from multiple replicates shown.

Because infected flies exhibit starvation-like effects on metabolite stores and insulin-pathway activity, we tested whether hyperactivity during infection was linked to infection-induced starvation signalling. Infection-induced anorexia has been observed in both mammals and insects (Adamo, 2005; Ayres and Schneider, 2009; Langhans, 2000; Wang et al., 2016) and hyperactivity is a known consequence of starvation in *D. melanogaster* (Keene et al., 2010; Lee and Park, 2004; Yang et al., 2015; Yu et al., 2016). We tested whether two infections we found were capable of increasing activity (*F. novicida* and *M. luteus*) also caused reduced food consumption. Surprisingly, we found that *F. novicida*-infected flies consumed 19.4% more food during infection compared to their mock controls and that food consumption was unaffected by infection with *M. luteus* (Figure 4C). Thus, infection-induced activity does not appear to be a by-product of anorexia.

Next, we tested the possibility that infection-induced increases in activity could be a product of endocrine signalling disruptions that mimicked the effects of starvation. In *D. melanogaster*, starvation increases activity via *adipokinetic hormone* (AKH) signalling in neurons (Lee and Park, 2004; Yu et al., 2016). We infected flies with a pan-neural reduction of the adiponkinetic hormone receptor (*nSyb*>*AkhR*-IR), a strategy previously shown to obliterate starvation-induced hyperactivity (Yu et al., 2016). Neuronal knockdown of *AkhR* did not affect activity during *F. novicida* infection, confirming that the infection-induced increase in activity here observed is distinct from a starvation response (Figure 4D). This finding is important because one of the proposed advantages of hyperactivity during starvation is greater resource acquisition from increased foraging. Thus, despite the failure of these infections to induce a starvation-like response via AKH, the results of these experiments are consistent with the idea that increased activity during *F. novicida* infection could lead to greater resource acquisition through increased feeding.

### Fat body derived *spz* contributes to *M. luteus* infection-induced activity

Bacterial peptidoglycan activates the Toll and IMD pathways, leading to the synthesis and secretion of antimicrobial peptides (AMPs) by the fat body in *D. melanogaster* (De Gregorio et al., 2002b; Lau et al., 2003; Lemaitre et al., 1995; Wang and Ligoxygakis, 2006). This production of AMPs contributes to the control of most bacterial infections. Since the Toll pathway is activated by Gram-positive bacteria (Hoffmann and Reichhart, 2002; Tanji and Ip, 2005), and we found that mutants of this pathway do not show an increase in activity during infection with the Gram-positive bacterium *M. luteus* (Figure 3B, C), we predicted that Toll signalling in the fat body played a role in infection-induced activity. To test this, we infected flies carrying fat body knockdowns of *spz* (*c564*>*spz*-IR), the circulating ligand that directly activates Toll; *MyD88* (*c564*>*MyD88*-IR), a key adaptor in the Toll pathway; and *Dif* (*c564*>*Dif*-IR), the primary *Toll*-activated NF-KB transcription factor in adult *Drosophila* (De Gregorio et al., 2002b; Valanne et al., 2011; Wang and Ligoxygakis, 2006). *spz* is synthesized and secreted as an inactive pro-protein where extracellular recognition factors initiate protease cascades that ultimately result in its proteolysis to produce active *spz*. This ligand binds and activates cell-surface Toll receptors (Alpar et al., 2018; An et al., 2010; Arnot et al., 2010; Valanne et al., 2011). As observed in the whole-body Toll signalling mutants, restricted knockdown of the Toll ligand *spz* to the fat body completely abolished the increase in activity observed during *M. luteus* infection (Figure 5A). Flies with *MyD88* or *Dif* knocked down in the fat body also exhibited reduced hyperactivity in response to *M. luteus* infection (Figure 5B, C). Importantly, the genetic control (*c564*>+) shows the expected increase of activity after bacterial infections (Figure 5 – figure supplement 1). These findings demonstrate that fat body-derived *spz* and fat body Toll pathway activity play a crucial role in the modulation of locomotor activity during infection.

**Figure 5.**
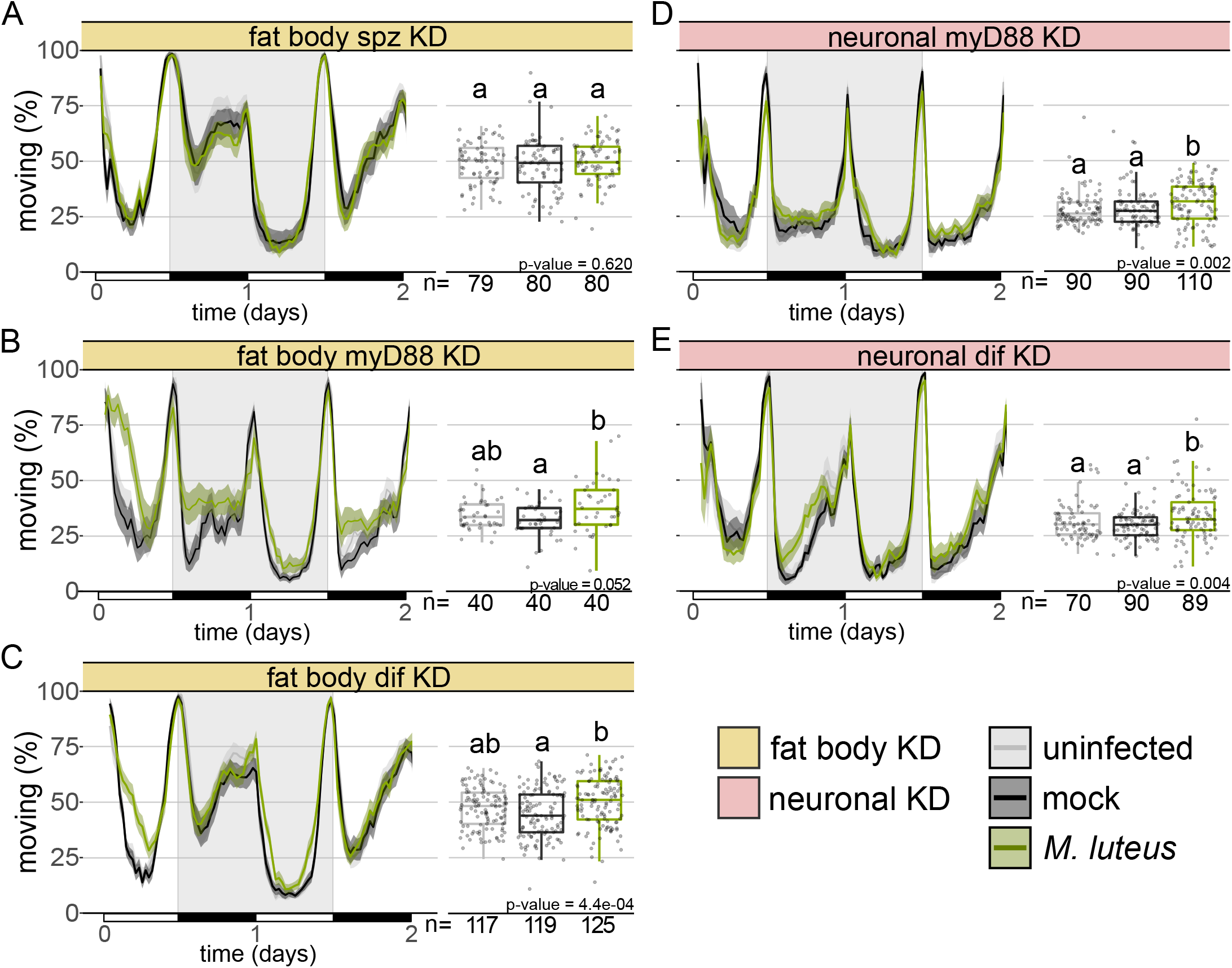
Fat body and neuronal Toll signalling is required for infection-induced activity during *Micrococcus luteus* infection. Ethogram showing percentage of flies moving over time with fat body **(A)** *spz* (w; *c564*> w; spz-IR) **(B)** *MyD88* (w; *c564*> w; *MyD88*-IR) and (C) *Dif* (w; *c564*> w; *Dif*-IR) knockdown. *Micrococcus luteus* infection had no effect on *spz*, nor *MyD88* fat body KD flies (*c564***>***spz***-IR**: Kruskal-Wallis chi-square = 0.96, df = 2, n = 239, p = 0.62; ***c564*>***MyD88***-IR**: Kruskal-Wallis chi-square = 5.92, df = 2, n = 120, p = 0.052). *Dif* fat body KD flies infected with *M. luteus* were significantly more active than mock-injected but not uninfected controls (***c564*>***Dif***-IR**: Kruskal-Wallis chi-square = 15.45, df = 2, n = 361, p = 4.4e-04; Dunn’s *post hoc*: mock|*M. luteus* = 2.6e-04, mock|uninfected = 0.077, uninfected| *M. luteus* = 0.053). Pan-neural **(A)** *MyD88* (*nSyb*>*MyD88*-IR) and **(B)** *Dif* knockdown (*nSyb*>*Dif*-IR) led to increased activity during infection (***nSyb*>***MyD88***-IR**: Kruskal-Wallis chi-square = 12.69, df = 2, n =290, p = 1.8e-03; Dunn’s *post hoc*: mock|*M. luteus* = 0.027, mock|uninfected = 0.304, uninfected| *M. luteus* = 1.7e-03; ***nSyb*>***Dif***-IR**: Kruskal-Wallis chi-square = 11.05, df = 2, n = 249, p = 3.9e-03; Dunn’s *post hoc*: mock|*M. luteus* = 5.7e-03, mock|uninfected = 0.68, uninfected| *M. luteus* = 0.018). Data from multiple replicates are shown.

### Neuronal KD of Toll signalling abrogates increased activity during *M. luteus* infection

Given that neither *spz* mutants nor *spz* fat body KD flies exhibit infection-induced activity, we thought that fat body-derived *spz* could be acting on another tissue to induce behavioural activity and decided to test whether neuronal knockdown of Toll signalling would also affect the activity phenotype. We infected flies with a pan-neural reduction of either *MyD88* (*nSyb*> *MyD88-*IR) or *Dif* (*nSyb*>*Dif*-IR). We found that knockdown of either *MyD88* or *Dif* reduced the increase in activity seen in *M. luteus* infected flies (Figure 5D, E), while the genetic control (*nSyb*>+) present the expected increase (Figure 5 – figure supplement 1). This effect was not complete—some increase in activity was still seen; this could reflect other mechanisms acting in parallel or it could be due to residual *MyD88/Dif* function in neurons. In either case, this finding lends support to our hypothesis that fat body-derived *spz* acts on other tissues – in this instance, neurons – to increase activity during *M. luteus* infection.

## Discussion

Here we show that bacterial infection in many cases leads to a marked increase in physical activity in *D. melanogaster*. This enhanced level of activity is mostly explained by an increase in walking (Figure 1). Though several bacteria induce activity upon infection in multiple fly lines with mutations in their immune response, we see pathogen/immune pathway specificity, as mutations in Toll signalling ablate activity-induction by some, but not all, bacteria. Finally, we demonstrate that neuronal Toll signalling plays a role in infection-induced changes in activity and propose that fat body-derived *spz* in required for this activation.

Immune activation has been shown to affect a range of behaviours and physiological functions including sleep, reproduction, cognition and metabolism (An and Waldman, 2016; Chambers et al., 2012; Dionne et al., 2006; Kobler et al., 2020; Kuo et al., 2010; Mallon et al., 2014; Shirasu-Hiza et al., 2007; Vincent and Sharp, 2014). Infection-induced changes in the host are often thought to be of benefit to either the host or the pathogen. Pathogen-mediated changes in host behaviour can lead to decreased survival, transmission and terminal host localisation (Adamo and Webster, 2013; Heil, 2016; Herbison et al., 2018; Lafferty and Shaw, 2013), whilst host-mediated changes during infection have been found to result in improved resistance and colony/conspecific protection (Boltaña et al., 2013; Sauer et al., 2019; Stroeymeyt et al., 2018; Ugelvig and Cremer, 2007). Interestingly, activity level within the first day of infection had a strong correlation with survival, though the direction of this relationship differed across bacterial strains (Figure 1E, Figure 2D, E). Whilst the reasons for the dissimilarity in the effect of activity on survival across these infections are unknown, what is consistent is the observation that early activity levels correlate with infection outcomes.

One well-documented change in host behaviour resulting from infection is anorexia (Adamo, 2005; Ayres and Schneider, 2009; Wang et al., 2016). Animals often decrease feeding in response to infection and this behaviour can be either harmful or beneficial to the host. In addition to infection-induced anorexia, *D. melanogaster* exhibits striking wasting phenotypes in a number of infections, characterized by decreased levels of glycogen and triglycerides (Chambers et al., 2012; Dionne et al., 2006; Vincent et al., 2020). Given the strong associations between infection and resource acquisition and utilization, one could imagine a scenario in which rather than infection directly leading to increased activity, instead, the metabolic dysregulation caused by infection signalled for increased activity as a means to acquire more food resource (Yang et al., 2015; Yu et al., 2016). Whilst both *M. luteus* and *F. novicida* infections led to strong wasting phenotypes, we found no evidence that starvation signalling contributed to increased activity, though increased activity was associated with greater food intake during *F. novicida* infection (Figure 4C).

While previous work found that bacteria-infected flies have poor quality sleep, as assessed through number of sleep bouts and bout duration (Shirasu-Hiza et al., 2007), these flies were not found to be more active than healthy controls. Infection with Gram-negative bacteria in *D. melanogaster* yields contrasting findings, showing that these infections can both reduce and increase sleep (Kuo et al., 2010; Kuo and Williams, 2014; Shirasu-Hiza et al., 2007). One study found that increased sleep lead to greater survival and bacterial clearance, but the flies studied were sleep deprived prior to infection, making it difficult to disambiguate the effect of the earlier sleep deprivation from – and on – subsequent infection and compensatory sleep. Interestingly, in that same study, flies in the control treatment – which were not sleep deprived – exhibited increased activity following infection (Kuo and Williams, 2014). We attribute the discrepancy between this study and previous work in observing a change in activity to differences between the annotative capabilities of the different activity monitoring systems used, specifically the greater spatial and temporal resolution afforded by the method employed here (Geissmann et al., 2017). Another potential explanation for the discrepancy is that while previous work evaluated activity levels immediately after the infection (Kuo et al., 2010; Kuo and Williams, 2014), we focus our attention on activity levels several hours or even days following initiation of the systemic infection. Thus, the temporal dynamics of the infection and the effects on behaviour may be intimately related.

Given the role of the fat body in the immune response, we predicted that pathogen recognition and subsequent activation of immune signalling pathways could contribute to the observed increase in activity; a supposition that was bolstered by the observation that both the occurrence and magnitude of the increased activity was positively correlated with the presence and number of bacteria. Disrupting the activity of the Toll pathway in the fat body phenocopied the ablation of activity observed in whole-body *spz* mutants, confirming that immune signalling in the fat body is vital during *M. luteus* infection. Whilst *spz* is secreted from tissues other than the fat body, these results suggest that the contribution of fat body derived *spz* is necessary. Our results leave open the possibility that fat body derived *spz* activates Toll signalling in neurons but further work is needed to confirm this interaction.

Intercellular signalling via cytokines has been shown to be vital to the induction of sickness behaviours (Dantzer, 2009, 2001; Davis and Raizen, 2017; Lasselin et al., 2020). Thus, immune detection in any of an organism’s organs has the potential to send signals to the brain that ultimately affect behaviour. One study found that in *D. melanogaster*, fat body derived *spz* was sufficient to induce sleep following infection (Kuo et al., 2010). Furthermore, a recent study showed that knocking down Toll signalling in the sleep-regulating R5 neurons suppressed the characteristic increase in sleep that is observed following sleep deprivation (Blum et al., 2021). Collectively these findings support a model where *spz* originating from the fat body, acts on a group of heretofore unidentified neurons to induce behavioural changes during infection. It follows that as pathogen load decreases, leading to less Toll signalling, we observe a corresponding extinction of infection-induced activity.

The role of increased activity during infection remains elusive but given that the behaviour appears to be activated via multiple pathways suggests that it serves a function rather than being the result of pleiotropy. Future work to discover other mechanisms involved has the potential to address the question of underlying function. Furthermore, the diversity of bacteria as well as the tools available to manipulate bacterial genomes, can be used to identify bacteria-derived signals that contribute to this response.

## Methods

### General experimental procedures

*w*^1118^ flies were used as wild-type flies throughout the study. A complete record of all other fly lines used in this study can be found in the supplementary information (Figure 2 supplementary table 1). For all experiments, male flies were collected following eclosion and kept in same-sex vials for 5 - 7 days in groups of 20. Thus, all experiments were conducted on flies between 5 and 8 days old. Flies were maintained on a standard diet composed of 10% w/v yeast, 8% w/v fructose, 2% w/v polenta, 0.8% w/v agar, supplemented with 0.075% w/v nipagin and 0.0825% vol propionic acid, at 25°C. Bacteria were grown from single colonies overnight at 37°C shaking with the exception of *L. monocytogenes* which was grown at 37°C without shaking. Each fly was injected with 50 nanolitres of bacteria diluted in PBS. Control flies were either injected with sterile PBS or were anaesthetized but otherwise unmanipulated, here referred to as mock controls and uninfected, respectively. Injections were carried out using a pulled-glass capillary needle and a Picospritzer injector system (Parker, New Hampshire, US). Following injection flies were kept at 29°C.

### Behavioural experiments

For all experiments, flies were sorted into glass tubes [70 mm × 5 mm × 3 mm (length × external diameter × internal diameter)] containing the same food used for rearing. After 2 days of acclimation under a regime of 12:12 Light:Dark (LD) condition in incubators set at 25°C animals were subject to either bacterial injection, mock injection, or anaesthetization (as above described), between zeitgeber time (ZT) 00 to ZT02 (the first 2h after lights ON) and transferred to fresh glass tubes containing our lab’s standard food as described above. Activity recordings were performed using ethoscopes (Geissmann et al., 2017) under 12:12 LD condition, 60% humidity at 29°C. Behavioural data analysis was performed in RStudio (RStudio Team, 2015) employing the Rethomics suit of packages (Geissmann et al., 2017). All behavioural assays were repeated at least twice with 20 – 60 flies/treatment/experiment. For lethal infections, behavioural data were analysed for the period between the first and final 12h of the assay; these windows of time were excluded as they encompass excessive noise due to awakening from anaesthesia/acclimation and mortality leading to declining sample sizes, respectively. For non-lethal infections, we analysed the 24h period following the initial 12h (t = 12h – 36h post infection), this time encompasses the duration of *M. luteus* infection after which live bacteria are no longer detected in flies.

### Bacterial quantification

Bacteria were quantified either via qPCR or plating. For plating, one fly was homogenised in 100μl of sterile ddH_2_O. Homogenates were serially diluted and plated onto LB agar plates where they incubated for 16-18h. Following incubation, the number of individual bacterial colonies observed on each plate was quantified and back-calculated to determine the number of CFUs present in each fly. For qPCR, one fly was homogenised in a 100μl of Tris-EDTA, 1% Proteinase K (NEB, P8107S) solution. Homogenates were incubated for 3h at 55°C followed by a ten-minute incubation at 95°C. Following incubation, we performed our qPCR protocol as outlined below to determine the number of bacterial colony forming units (CFU). All quantifications were repeated at least twice with 8-16 samples/treatment/experiment.

### Measurement of triglycerides

Triglycerides were measured using thin layer chromatography (TLC) assays as described elsewhere (Al-Anzi et al., 2009). Briefly, each sample consisted of 10 flies; flies were placed in microcentrifuge tubes and stored at −80°C until the time of analysis. To perform the TLC assay, samples were removed from the −80°C freezer and spun down (3 min at 13,000 rpm at 4°C) in 100μl of a chloroform (3) : methanol (1) solution. Flies were then homogenised and subject to a further ‘quick spin’. Standards were generated using lard dissolved in the same chloroform : methanol solution. We loaded 2μl of each standard and 20μl of each sample onto a silica gel glass plate (Millipore). Plates were then placed into a chamber pre-loaded with solvent (hexane (4) : ethyl ether (1)) and left to run until the solvent could be visualised 1cm prior to the edge of the plate. Plates were then removed from the chamber, allowed to dry, and stained with CAM solution. Plates were baked at 80°C for 15-25min and imaged using a scanner. Analysis was conducted in ImageJ using the Gels Analysis tool. This assay was repeated at least twice with four samples/treatment/experiment.

### Measurement of carbohydrates (glucose + trehalose and glycogen)

Each sample contained three flies that were homogenised in 75μl of TE + 0.1% Triton X-100 (Sigma Aldrich). Samples were incubated for 20 min at 75°C and stored at −80°C. Prior to the assay, samples were incubated for 5 min at 65°C. Following incubation, 10μl from each sample was loaded into four wells of a 96-well plate. Each well was designated to serve as a measurement for either: control (10μl sample + 190μl H_2_0), glucose (10μl sample + 190μl glucose reagent (Sentinel Diagnostics)), trehalose (10μl sample + 190μl glucose reagent + trehalase (Sigma Aldrich)), or glycogen (10μl sample + 190μl glucose reagent + amyloglucosidase (Sigma Aldrich)). A standard curve was generated by serially diluting a glucose sample of known concentration and adding 190μl of glucose reagent to 10μl of each standard. Standards were always run at the same time and in the same plate as samples. Plates were incubated for 1h at 37°C following which the absorbance for each well at 492 nm was determined using a plate reader. This assay was repeated at least twice with four samples/treatment/experiment.

### Western Blots

Each sample contained three flies that were homogenised directly in 75μl 2x Laemmli SDS-PAGE buffer. The primary antibodies used were anti-phospho-Akt (Cell Signalling Technologies 4054, used at 1:1000), anti-total-Akt (Cell Signalling Technologies 4691, 1:1000), and anti-α-tubulin (Developmental Studies Hybridoma Bank 12G10, 1:5000). The secondary antibodies used were anti-rabbit IgG (Cell Signalling Technologies 7074, 1:5000) and anti-mouse IgG (Cell Signalling Technologies 7076, 1:10,000). The chemiluminescent substrate used was SuperSignalTM West Pico PLUS (Thermo Scientific 34580). Blots were imaged using a Fuji LAS-3000 luminescent image analyser and images analysed in ImageJ.

### Feeding

For each sample, eight flies were placed into a 50mL Falcon tube with a lid containing our standard food (described above) with the addition of the food dye Erioglaucine sodium salt, 1% w/vol (Alfa Aesar) and left for 30 or 80h. To determine the amount of ingested food, flies were homogenised in 200μL of Tris-EDTA 0.1% Triton-X. Following homogenisation samples were spun down (20 minutes at 13 000 rpm, RT) and 100μL of the supernatant removed (this contained predominantly suspended triglyceride). We then added 300μL of Tris-EDTA 0.1% Triton-X to the sample and spun for 10 minutes at 13 000 rpm, RT. 200μL of each suspension was placed into a 96-well plate. To determine the amount of excreted food, 1mL of Tris-EDTA 0.1% Triton-X was added to each Falcon tube and the tube briefly vortexed. Following vortex, tubes were placed on a roller for 5 minutes and then subject to a ‘quick-spin’. 200μL was taken from each tube and placed into a 96-well plate. 96-well plates were read at 620nm and normalized to the mean value of the uninfected controls. Data are presented as a combination of excreted (Falcon tube) and ingested (fly homogenate) values. This assay was repeated at least twice with four samples/treatment/experiment.

### Statistical analysis

Data were analysed in R Studio with R versions 3.5.3 and 3.6.3 (RStudio Team, 2015). Behavioural data were analysed using the Rethomics package (Geissmann et al., 2017); for all other assays, we first tested for normality of data which dictated whether an ANOVA, t-test, Kruskal-Wallis analysis of variance, or Mann-Whitney U test was used to calculate differences between treatments. When appropriate, we performed *post hoc* Tukey, Nemenyi or Dunn analyses to identify specific differences between treatments. All assays were repeated at least twice with sample sizes as indicated within the reported statistics.

## Acknowledgements

We are indebted to members of the Dionne and Gilestro labs for critical discussion. Q. Geissmann provided invaluable feedback and support in the behavioural quantification. This work was funded by Wellcome Trust Investigator Award 207467/Z/17/Z, MRC Research Grants MR/R00997X/1 and MR/L018802/2, and BBSRC Research Grant BB/L020122/2 to MSD, and BBSRC Research Grant BB/R018839/1 to GG. KK was supported by a DFG fellowship. EJB was supported by the People Programme (Marie Curie Actions) of European Union’s Eighth Framework Programme H2020 under REA grant agreement 705930.

**The authors declare that they have no conflict of interest**.

---

**Figure 1 - Supplementary Figure 1. Quantifying engagement in specific behaviours.** Ethogram showing percentage of infected wild-type males **(A)** walking, **(B)** engaging in micro-movements (e.g. feeding and grooming) and **(C)** sleeping. Boxplots show the quantification of **(D)** total distance covered and **(E)** the total number of times that the flies crossed the middle of the housing tube as a proportion of the time (in seconds) spent awake. Uninfected and mock controls are represented by grey and black tracings, respectively. Infected flies are in blue. **Distance covered when active** (Kruskal-Wallis chi-square = 6.496, df = 2, n = 419, p = 0.039; Dunn’s *post hoc*: mock|*F. novicida* = 0.033, mock |uninfected = 0.163, uninfected|*F*. *novicida* = 0.460) and **midline crosses when active** (Kruskal-Wallis chi-square = 5.453, df = 2, n = 419, p = 0.065; Dunn’s *post hoc*: mock|*F. novicida* = 0.064, mock |uninfected = 0.330, uninfected|*F*. *novicida* = 0.339) were not impacted by the infection. **(F)** Activity level throughout *F. novicida* infection was not correlated with survival (Pearson’s correlation, *r* = 0.313; t = 3.31, df = 101, p = 0.001) and (G) activity levels on day 1 are positively correlated with total activity (Pearson’s correlation, *r* = 0.744; t = 11.2, df = 101, p = 2.2e-16). Data from multiple replicates are shown.

**Figure 2 - Supplementary Figure 1. Not all bacteria induce activity in wild-type flies**. Ethograms showing percentage of flies moving over time. Uninfected and mock controls are represented by grey and black tracings, respectively. Infected flies are in orange. Neither of these extracellular/Gram-negative bacteria, *Escherichia coli* and *Enterobacter cloacae* induced activity (***E. coli***: Kruskal-Wallis chi-square = 3.699, df = 2, n = 300, p = 0.16; ***E. cloacae***: Kruskal-Wallis chi-square = 1.516, df = 2, n = 229, p = 0.47). Data from multiple replicates are shown.

**Figure 2 - Supplementary Figure 2. *Francisella novicida* infection increases activity in several immune and locomotor mutants.** Boxplots showing the percentage of flies moving over time. Uninfected and mock controls are represented by grey and black tracings, respectively. Infected flies are in blue. Previously characterized phenotypes and statistics of studied mutants can be found in Figure 2 – Supplementary Tables 1 and 2. Data from multiple replicates are shown.

**Figure 2 –figure supplement table 1.** Phenotypes of activity mutants tested.

| Gene | Published phenotype |
| --- | --- |
| yw | Alternative ‘control’ line; abnormal colour (yellow) & male courtship behaviour |
| *tak1*<sup>1</sup> | Highly susceptible to Gram negative bacteria |
| *upd2*<sup>Δ</sup> | Defective immune response |
| *pdf*<sup>01</sup> | Locomotor & circadian behaviour defective |
| *dop1R2*<sup>MB05108</sup> | Hypoactive |
| *dopR1*<sup>f2676</sup> | Hyperactive |
| *iav*<sup>3621</sup> | Locomotor behaviour defective |

\*references which cite each phenotype can be found via allele pages on FlyBase.

**Figure 2 –figure supplement table 2.** Statistics from activity mutant assays.

| Genotype | Kruskal-Wallis Test Statistics | Uninfected\|PBS | Uninfected\|*F. novicida* | PBS\|*F. novicida* |
| --- | --- | --- | --- | --- |
| y,w; | X<sup>2</sup>= 12.456, df= 2, n= 220, p= 1.8e-03 | p = 0.14 | p = 0.041 | p = 1.3e-03 |
| *tak1*<sup>1</sup>/FM7h;; | X<sup>2</sup>= 32.963, df= 2, n= 258, p= 6.9e-08 | p = 0.77 | p = 3.6e-07 | p = 4.1e-06 |
| w *upd2*<sup>Δ</sup>;; | X<sup>2</sup>= 66.744, df= 2, n= 237, p= 3.2e-15 | p = 0.26 | p = 1.4e-10 | p = 1.5e-13 |
| w;;*pdf*<sup>01</sup> | X<sup>2</sup>= 13.525, df= 2, n= 237, p= 1.2e-03 | p = 0.27 | p = 0.02 | p = 1e-03 |
| *dop1R2*<sup>MB05108</sup> | X<sup>2</sup>= 15.524, df= 2, n= 90, p= 4.3e-04 | p = 0.94 | p = 1.1e-03 | p = 1.6e-03 |
| *dopR1*<sup>f2676</sup> | X<sup>2</sup>= 44.393, df= 2, n=130, p= 2.3e-10 | p = 0.81 | p = 1.4e-08 | p = 1.1e-08 |
| *iav*<sup>3621</sup>/FM7h | X<sup>2</sup>= 32.974, df= 2, n= 158, p= 6.9e-08 | p = 0.36 | p = 1.6e-05 | p = 3.8e-07 |

**Figure 2 –Supplementary Figure 3. *Francisella novicida* and *Micrococcus luteus* differ in lethality.** In all plots, grey and black tracings represent uninfected and mock controls, respectively. *F. novicida*, *Micrococcus luteus*, *Listeria monocytogenes* and *Staphylococcus aureus* infections are shown in blue, green, orange and yellow, respectively. *Francisella novicida* infection was lethal in all four genotypes. Infection with *M. luteus* did not result in more lethality than either uninfected or mock controls, while *L. monocytogenes* and *S. aureus* both lead to decreased survival. Median survival is indicated by dotted lines intersecting the y and x axes at 50% survival and time (in days), respectively. Survival was calculated at the same time as activity data and thus have the same sample size as indicated elsewhere. Data from multiple replicates are shown. **Quantification of** *M. luteus* load markers (as indicated) represent means and whiskers represent SE. Initial inoculum consisted of ~ 10^3^ colony forming units (CFUs). Within 30h bacterial numbers decreased to near-undetectable levels (average of 28-40 CFUs/fly). **Quantification of *F. novicida*.** Bacterial numbers increase over the course of infection. All genotypes were injected with the same initial dose (t = 0; ~1700 CFUs). The last measured timepoint was 24h prior to the onset of death for each genotype; this was 72h for all genotypes except *imd^10191^* which was 48h. Genotypes are represented by marker style and line colour as indicated inset. Markers indicate means and whiskers represent SE. Bacterial quantifications were repeated at least twice, n = 16-22 flies/genotype/timepoint; data from all replicates are shown.

**Figure 3 –Supplementary Figure 1. Mutants of Toll and IMD pathways exhibit increased locomotor activity during *Francisella novicida* infection.** Ethogram showing percentage of **(A)** *imd*^10191^ **(B)** *spz*^eGFP^ and **(C)** *imd*^10191^; *spz*^eGFP^, flies moving over time. *Francisella novicida* infected animals moved significantly more than both the uninfected and mock controls (***imd***: Kruskal-Wallis chi-square = 111.32, df = 2, n = 482, p = 2.2e-16; Dunn’s *post hoc*: mock|*F. novicida* = 1.3e-20, mock|uninfected = 0.38, uninfected|*F. novicida* = 1.4e-18; ***spz***^eGFP^: Kruskal-Wallis chi-square = 59.59, df = 2, n = 220, p = 1.1e-13; Dunn’s *post hoc*: mock|*F. novicida* = 1.7e-12, mock|uninfected = 0.36, uninfected|*F. novicida* = 1.6e-09; ***imd;spz***^eGFP^: Kruskal-Wallis chi-square = 52.594, df = 2, n = 195, p = 3.8e-12; Dunn’s *post hoc*: mock|*F. novicida* = 5.6e-09, mock|uninfected = 0.45, uninfected|*F. novicida* = 1.9e-10). Data from multiple replicates are shown.

**Figure 5 –Supplementary Figure 1. Fat body and pan-neural driver controls exhibit increased locomotor activity during infection. (A)** Ethogram showing percentage of *c564*>**+** flies moving over time. *Francisella novicida-*infected animals moved significantly more than both the uninfected and mock controls (Kruskal-Wallis chi-square = 8.41, df = 2, n = 128, p = 0.015; Dunn’s *post hoc*: mock|*F. novicida* = 0.049, mock|uninfected = 0.41, uninfected|*F. novicida* = 0.015). *Micrococcus luteus*-infected animals moved significantly more than uninfected controls but no more than mock controls (Kruskal-Wallis chi-square = 7.64, df = 2, n = 136, p = 0.022; Dunn’s *post hoc*: mock|*M. luteus* = 0.091, mock|uninfected = 0.35, uninfected|*M. luteus* = 0.022). **(B)** *Francisella novicida-*infected *nSyb>***+** flies moved significantly more than both the uninfected and mock controls (Kruskal-Wallis chi-square = 46.38, df = 2, n = 178, p = 8.5e-11; Dunn’s *post hoc*: mock|*F. novicida* = 3.6e-08, mock|uninfected = 0.56, uninfected|*F. novicida* = 1.8e-09). *Micrococcus luteus*-infected animals moved significantly more than uninfected controls but no more than mock controls (Kruskal-Wallis chi-square = 7.35, df = 2, n = 179, p = 0.025; Dunn’s *post hoc*: mock|*M. luteus* = 0.19, mock|uninfected = 0.24, uninfected|*M. luteus* = 0.021). *Francisella novicida* and *Micrococcus luteus* data were analysed over 0.5d - 3.5d and 0.5d - 1.5d, respectively. Data from multiple replicates are shown.

